# Physiological noise optimizes multiplexed coding of vibrotactile-like signals in somatosensory cortex

**DOI:** 10.1101/2021.09.11.459897

**Authors:** Mohammad Amin Kamaleddin, Aaron Shifman, Daniel MW Sigal, Steven A Prescott

## Abstract

Neurons can use different aspects of their spiking to simultaneously represent (multiplex) different features of a stimulus. For example, some pyramidal neurons in primary somatosensory cortex (S1) use the rate and timing of their spikes to respectively encode the intensity and frequency of vibrotactile stimuli. Doing so has several requirements. Because they fire at low rates, pyramidal neurons cannot entrain 1:1 with high-frequency (100-600 Hz) inputs and instead must skip (*i*.*e*. not respond to) some stimulus cycles. The proportion of skipped cycles must vary inversely with stimulus intensity for firing rate to encode stimulus intensity. Spikes must phase lock to the stimulus for spike times (intervals) to encode stimulus frequency but, in addition, skipping must occur irregularly to avoid aliasing. Using simulations and *in vitro* experiments in which S1 pyramidal neurons were stimulated with inputs emulating those induced by vibrotactile stimuli, we show that fewer cycles are skipped as stimulus intensity increases, as required for rate coding, and that physiological noise induces irregular skipping without disrupting phase locking, as required for temporal coding. This occurs because the reliability and precision of spikes evoked by small-amplitude, fast-onset signals are differentially sensitive to noise. Simulations confirmed that differences in stimulus intensity and frequency can be well discriminated based on differences in spike rate or timing, respectively, but only in the presence of noise. Our results show that multiplexed coding by S1 pyramidal neurons is facilitated rather than degraded by physiological levels of noise. In fact, multiplexing is optimal under physiologically noisy conditions.

## INTRODUCTION

Neurons can encode information using the rate or timing of their action potentials (spikes). The distinction between rate and temporal coding depends on the temporal resolution used to decode responses (Rieke et al., 1997), which itself depends on the integrative properties of downstream neurons – integrators and coincidence detectors have long and short integration times, respectively (König et al., 1996; Ratté et al., 2013). Using a wide time window (low resolution) means that each window can contain multiple spikes, enabling the spike count to carry information. Conversely, using a narrow window (high resolution) ensures that each window contains no more than one spike, with most containing none, thus revealing whether the intervals between spikes carry information. The appropriate window size depends on the signal.

Rate and temporal coding are not mutually exclusive. These two coding schemes can co-exist if information about different stimulus features is encoded independently (multiplexed) and if decoders can extract (demultiplex) that information. For instance, neurons in S1 can encode the intensity of tactile stimuli and when intensity changes using the rate of asynchronous spikes and the timing of transiently synchronized spikes, respectively (Lankarany et al., 2019); in other words, each stimulus feature is represented by separate spikes distinguished by their degree of synchronization. Harvey et al. (2013) described a different form of multiplexing in which the amplitude and frequency of periodic (vibrotactile) stimuli are encoded by the rate and timing of the same spikes. The current study focusses on the latter scenario.

Grappling with similar issues in the auditory system, Wever and Brey (1930) proposed the volley theory nearly a century ago to explain how sound intensity and pitch could be represented by the rate and timing of spikes in the auditory nerve. They posited that spikes occur at a preferred phase of the stimulus (*i*.*e*. phase lock), but since auditory afferents cannot (because of their refractory period) fire at the kHz frequencies associated with high-pitch sounds, they reasoned that only a fraction of all stimulus cycles evoke a spike. After generating a spike, a neuron does not fire during subsequent stimulus cycles until its refractoriness has waned. Firing rate encodes stimulus intensity (amplitude) because the stronger the stimulus, the fewer cycles are skipped before the neuron fires again. Spike timing encodes stimulus frequency (pitch) because, if the refractoriness of different neurons wears off at different rates, different neurons respond to different stimulus cycles, allowing the population as a whole to entrain to the stimulus.

Volley theory is directly relevant for the responses to vibrotactile stimulation reported by Harvey et al. (2013) in S1 neurons, but unlike Wever and Bray’s theory, they found that individual neurons skipped a variable number of stimulus cycles after each spike. Irregular skipping could result from noise, like in stochastic resonance (Longtin, 1993; McDonnell and Ward, 2011). Noise can originate from many sources (Faisal et al., 2008), including the stochastic opening and closing of ion channels (White et al., 2000) and background synaptic input, which is especially prominent in the intact cortex (Destexhe and Paré, 1999; Destexhe et al., 2003). But since noise can severely disrupt spiking timing (London et al., 2010), it is unclear whether noise could induce irregular skipping without disrupting phase locking. Indeed, precisely timed spikes often occur very reliably, as in synfire chains (Abeles, 1991), but multiplexing requires that precisely timed spikes have a modulatable rate.

We therefore sought to identify under what conditions rate and temporal coding co-exist by elucidating how the rate and timing of spikes can be independently controlled. Through simulations and *in vitro* experiments, we found that both coding schemes benefit from physiological levels of noise because such noise affects the probability (reliability) more than the timing (precision) of spikes driven by small-amplitude, fast-onset signals. Our results demonstrate that multiplexed coding of vibrotactile stimuli is not only possible under realistically noisy conditions, it actually benefits from that noise.

## RESULTS

### Multiplexed coding of sinusoidal input by a model neuron

If a neuron spikes once on each cycle of a periodic stimulus (*i*.*e*. entrains 1:1), then firing rate will scale with stimulus frequency (**Fig. 1A**). This entails spiking at rates much higher than pyramidal neurons can maintain and also precludes rate coding of stimulus amplitude. To explore multiplexed coding of periodic signals at physiological firing rates, we first conducted a series of simulations in an adaptive exponential (AdEx) integrate-and-fire model with parameters adjusted so to yield evoked firing rates <50 spikes/s.

**Figure 1.**
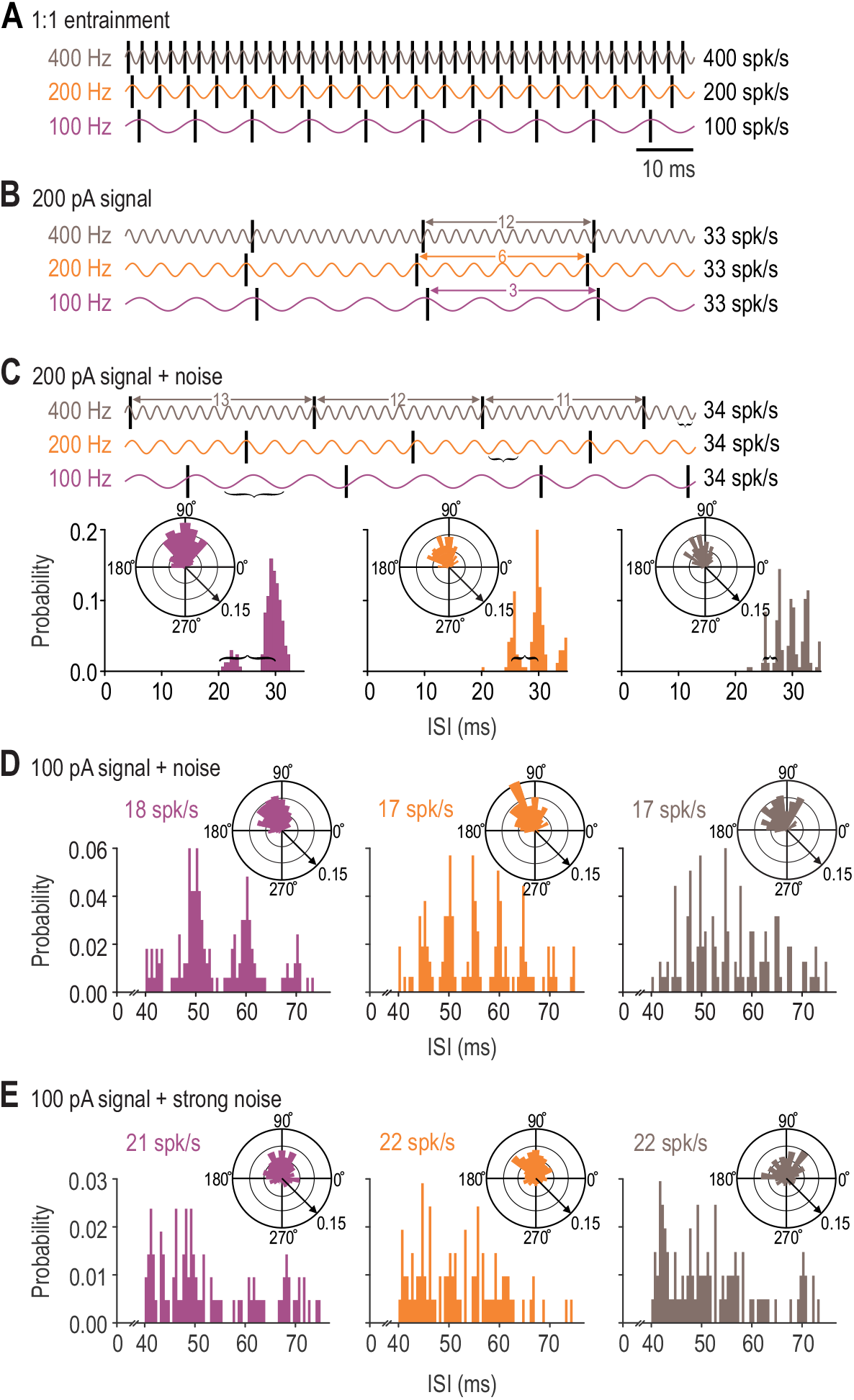
Multiplexed coding of stimulus amplitude and frequency in a model neuron. **(A)** Cartoon rasters overlaid on 100, 200 or 400 Hz sinusoidal stimulation (purple, orange and grey, respectively) to illustrate 1:1 entrainment of spiking with the stimulus. Neither our AdEx model nor real pyramidal neurons fire at such high rates. **(B)** Rasters from the AdEx model injected with 200 pA peak-to-peak sinusoidal current. The model responded to all stimulus frequencies with 33 spikes/s by spiking on every third, sixth, or 12^th^ cycle (see double-headed arrows) during 100, 200 and 400 Hz stimulation. **(C)** Responses to same stimulus as in **B** but with added noise. Polar plots show that spikes phase lock (*i*.*e*. occur at a relatively consistent phase of the stimulus cycle) despite noise. Rather than spiking after a consistent number of stimulus cycles like in **B**, noise caused irregular skipping, resulting in a multimodal ISI distribution with peaks separated by the cycle period (brackets). Average firing rate was 34 spikes/s for all stimulus frequencies. **(D)** Responses to 100 pA peak-to-peak sinusoidal current with noise like in **C**. The ISI distribution remained multimodal but shifted to the right, consistent with the reduced firing rate of 17-18 spikes/s. Please note difference in scaling of x-axis. **(E)** Responses to same stimulus as in **D** but with stronger noise. Phase locking was diminished (see polar plots) and peaks on the ISI distribution were obscured. Rings on each polar plot represent the probability of spiking; arrow indicates scale.

When a sine wave with a peak-to-peak amplitude of 200 pA was applied to the model neuron, spikes occurred on only a subset of cycles to produce a physiologically realistic firing rate of 33 spikes/s (**Fig. 1B**). A spike occurred on every third, sixth, or 12^th^ stimulus cycle to yield the same firing rate during 100, 200, and 400 Hz stimulation, respectively; in other words, more cycles were skipped at higher stimulus frequencies. But if skipping causes all spikes to occur at the same interval (33 ms in this case), equivalent spiking could arise from 1:1 entrainment during 33 Hz stimulation or any integer multiple thereof (*e*.*g*. 1:2 entrainment for 66 Hz stimulation, 1:3 for 100 Hz stimulation, etc.). Stimulus frequency cannot be unambiguously decoded from such responses.

The ambiguous representation of stimulus frequency in Figure 1B is due to patterning introduced by the regularity of skipping. This aliasing can be avoided by skipping stimulus cycles irregularly, as is likely to occur under noisy conditions (see Introduction). As expected, when noise approximating background synaptic activity was added to the simulation, the model neuron still responded to 200 pA sinusoidal current with an average rate of ∼33 spikes/s, but stimulus cycles were now skipped irregularly, resulting in a multimodal distribution of interspike intervals (ISIs) (**Fig. 1C**). The spacing between histogram peaks corresponds to the cycle period of 10, 5, and 2.5 ms for 100, 200, and 400 Hz stimulation, respectively; in other words, spikes occur at intervals corresponding to different integer multiples of the cycle period. Even though spikes never occur at the cycle period itself (which is shorter than the neuron’s refractory period), the fundamental period can be unambiguously inferred from the multimodal ISI distribution.

When a weaker stimulus was tested under equivalently noisy conditions, the ISI histogram still exhibited several evenly spaced peaks but the overall distribution shifted to a longer average ISI (**Fig. 1D**), consistent with rate coding of stimulus amplitude – more stimulus cycles were skipped at lower stimulus amplitudes. In other words, the ISI histogram’s shape (*i*.*e*. spacing between peaks) contains information about stimulus frequency whereas the histogram’s central tendency (*i*.*e*. average ISI) contains information about stimulus amplitude. Plotting ISI histograms using narrow or wide bins helps emphasize each type of information (see Fig. 2C). Notably, spikes must occur at a consistent phase of the stimulus (*i*.*e*. phase lock; see polar plots on Fig 1C,D) to produce a multimodal ISI distribution. Strong noise diminishes phase locking, smearing those peaks (**Fig. 1E**) and compromising temporal coding.

**Figure 2.**
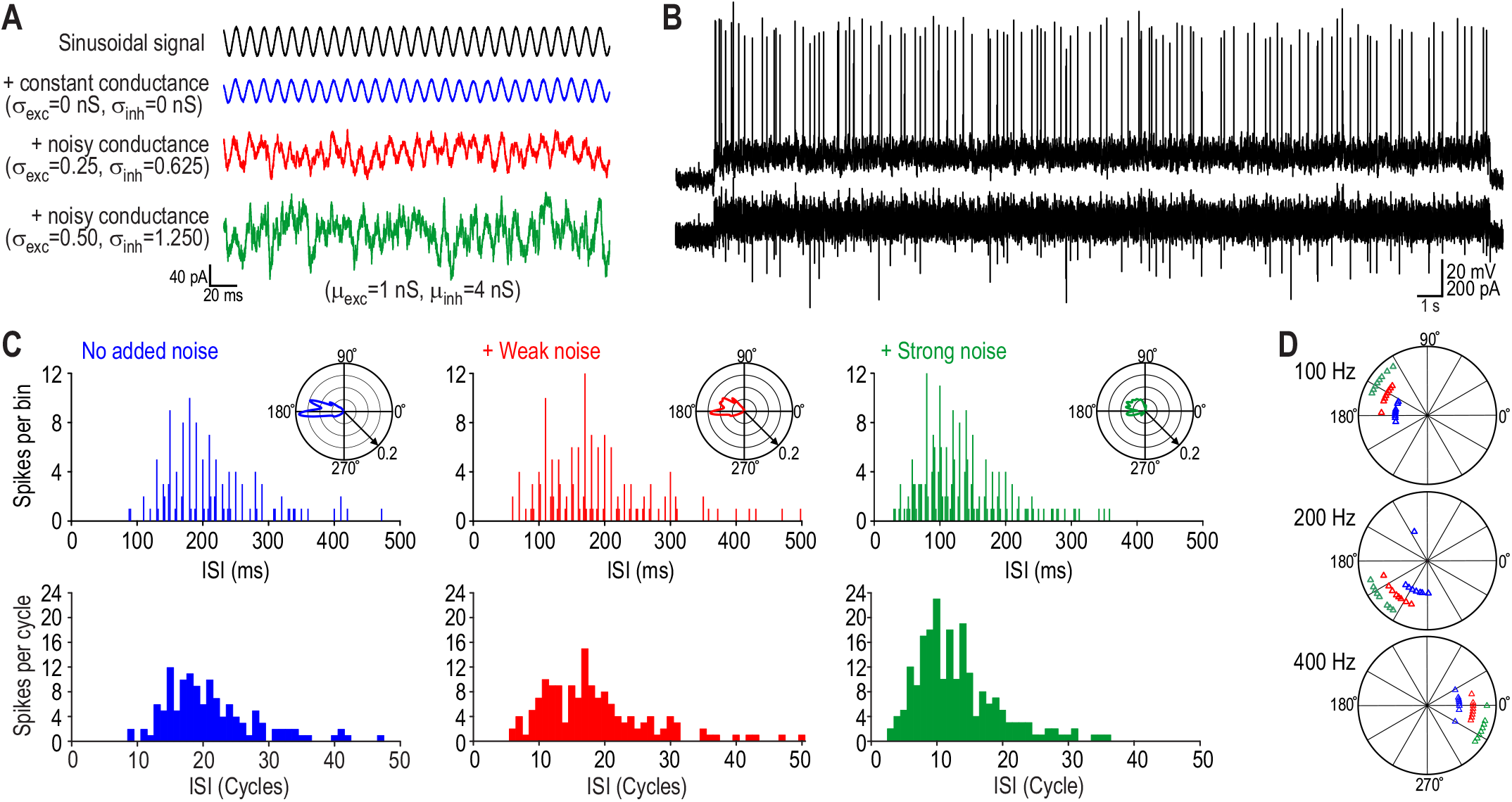
Response of a typical pyramidal neuron to sinusoidal current injection with and without added noise. **(A)** Periodic stimuli were applied as sinusoidal current (50 or 100 pA at 100, 200 or 400 Hz) with a DC offset (see Methods). Dynamic clamp was used to apply a constant conductance (blue) or fluctuating conductance approximating weak (red) or strong (green) noise. **(B)** Sample response from a pyramidal neuron stimulated with a 100 pA, 100 Hz sine wave plus strong noise. Dynamic clamp starts before the 30 s-long stimulus and continues throughout. **(C)** Responses from the same neuron to 100 pA, 100 Hz stimulation at three noise levels. Top histograms (with bin widths set to 2 ms = 1/5 of cycle period) exhibit multiple peaks separated by the cycle period of 10 ms. Bottom histograms (with bin width set to 10 ms) emphasize the number of cycles between each spike. Increased firing rate with increasing noise is reflected by a leftward shift in the ISI distribution. **(D)** Median phase from the 8 neurons with the highest firing rate in which all combinations of stimulus frequency and noise level were tested.

These results show that rate and temporal coding can co-exist in a model neuron, but only if there is neither too much nor too little noise. An important question, therefore, is whether real pyramidal neurons can support the same multiplexed coding under conditions that exist *in vivo*.

### Multiplexed coding of sinusoidal input by pyramidal neurons

We proceeded to test L2/3 pyramidal neurons in slices of S1 prepared from adult mice. Periodic signals were applied as sinusoidal current (or as synaptic trains in later experiments) while virtual background synaptic input (noise) was applied using dynamic clamp (**Fig. 2A**). Real synaptic input was blocked pharmacologically. Non-fluctuating conductance was applied in the *no added noise* condition so that the average additional conductance due to background input was equivalent across noise levels, reducing input resistance by ∼70%, consistent with the difference between *in vitro* and *in vivo* recordings from L2/3 pyramidal neurons in S1 (Fernandez et al., 2018). The *weak noise* condition, which produced membrane potential fluctuations with a standard deviation of 2-2.5 mV, is our best estimate of the noise experienced by L2/3 pyramidal neurons (Fernandez et al., 2018). **Figure 2B** shows a typical response to a 30 s-long stimulus with strong noise. The injected current fluctuates during spikes because the driving force of dynamic clamp conductances depends on membrane potential. To avoid time-dependent changes in firing rate due to adaptation, only spikes from the last 28 seconds of each response were used for analysis.

**Figure 2C** shows analysis from a typical neuron responding to 100 Hz, 100 pA sinusoidal current. Polar plots show that spikes phase lock to the periodic stimulus, though less so with increasing noise. Histograms constructed with narrow bins (top) exhibit multiple peaks separated by the cycle period of 10 ms, consistent with irregular skipping. Those peaks are not evident on histograms constructed with wide bins (bottom), which instead highlight the number of stimulus cycles between each spike. Notably, irregular skipping was evident across all noise levels, even when no synaptic noise was added. Unlike simulations, which are truly noise-free until noise is added to the model neuron, noise intrinsic to real neurons (*e*.*g*. channel noise) can evidently induce irregular skipping. The preferred phase was consistent across neurons (**Fig. 2D**).

Next, we asked how stimulus parameters and noise level affect phase locking. Given that phase preference is consistent across neurons (see Fig. 2D), data from all neurons that responded to stimulation with firing rates >1 spike/s were pooled to assess phase locking (**Fig. 3A**). Higher frequency stimulation was associated with spikes occurring later in the cycle and with a broader phase distribution, which suggests more variable spike timing (jitter), but both effects are due to the shorter period associated with high-frequency stimulation. Re-plotting histograms against time (**Fig. 3B**), rather than phase, reveals that jitter is lowest during high-frequency stimulation. For 400 Hz stimulation, spike latencies of 2.37 ± 0.52 (mean ± SD) relative to the cycle period of 2.5 ms mean that some spikes occur as late as the start of the next cycle, whereas latencies of 3.20 ± 0.77 and 4.61 ± 1.28 lie within the cycle periods of 5 and 10 ms for 200 and 100 Hz stimulation, respectively. For the values above, spike times were pooled across stimulus intensities and noise levels. Plotting mean latency against jitter for each combination of stimulus frequency, stimulus amplitude, and noise level (**Fig. 3C**) shows that jitter depends on noise level and stimulus amplitude, whereas mean latency depends on stimulus frequency.

**Figure 3.**
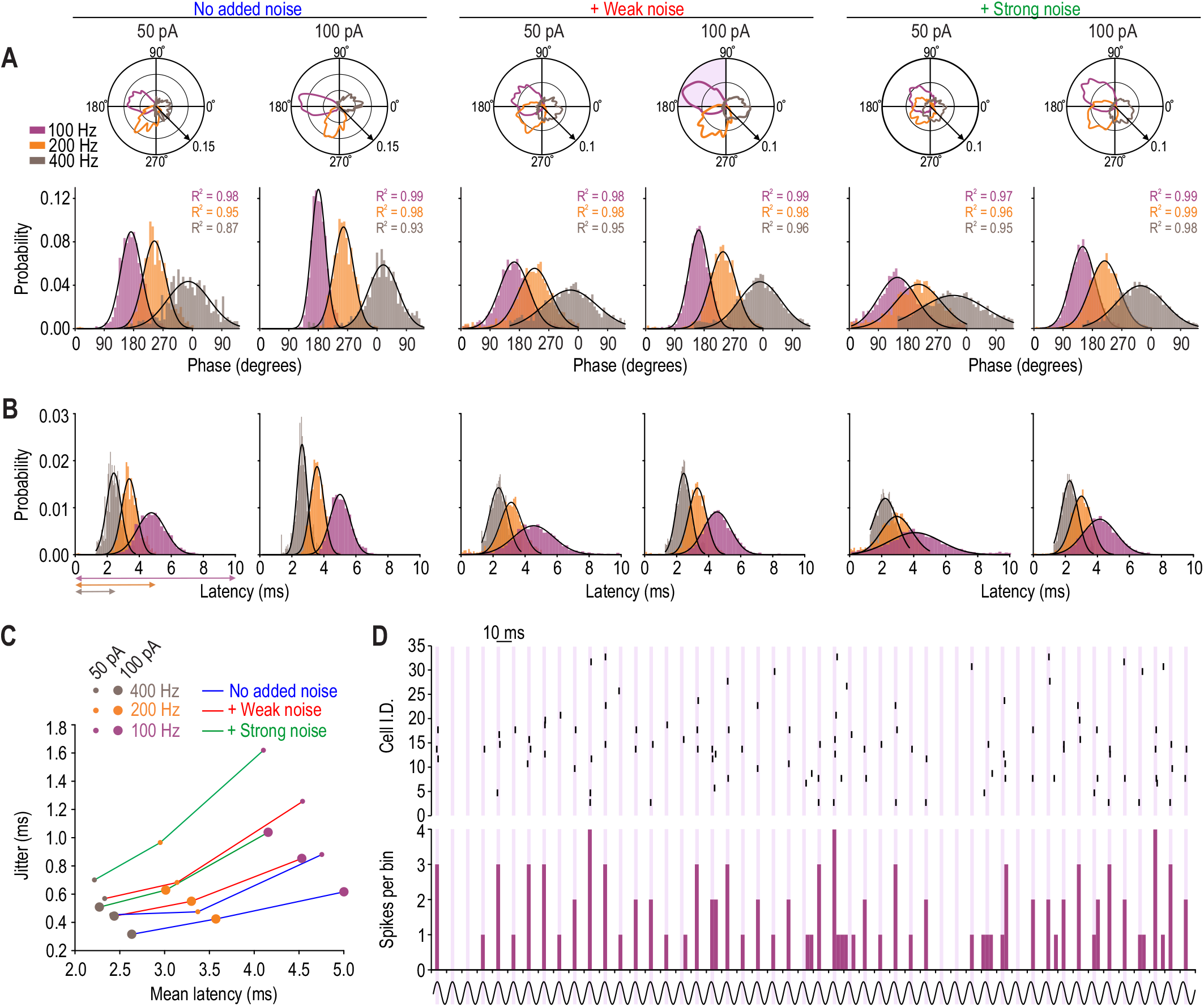
Spikes phase lock to sinusoidal input despite noise. **(A)** Polar plots (top) and histograms (bottom) show distribution of spike times relative to stimulus phase. Data for each condition were aggregated across all neurons (*n* = 32) and fit with a Gaussian curve (black); coefficients of determination (*R*^2^) are reported on the graphs. The polar plot with shading in its top left corner corresponds to conditions in panel **D**; phase preference between 90- 180° corresponds to latency of 2.5 - 5 ms. Spikes evoked by higher frequency inputs occurred later in the stimulus cycle, with many spikes evoked by 400 Hz input occurring near the start of the subsequent cycle. Distributions broadened when noise was increased, stimulus amplitude was reduced, or stimulus frequency was increased. **(B)** Data from **A** plotted against time rather than phase. Spikes evoked by 400 Hz input occurred with the shortest latency and with the narrowest temporal distribution (jitter). Arrows underneath leftmost panel indicate stimulus period. **(C)** Mean spike latency plotted against jitter for each combination of stimulus frequency, stimulus amplitude and noise level. **(D)** Rasters (top) and firing rate histogram (bottom) constructed using a 2.5 ms bin for 100 Hz, 100 pA stimulation plus weak noise. Most spikes occurred between 2.5 and 5 ms after the start of each stimulus cycle (purple shading); compare with shaded polar plot in **A**. Each neuron responded to relatively few stimulus cycles but each cycle typically evoked spikes in 1-3 neurons (out of 32) resulting in clearly spaced (temporally precise) peaks on the firing rate histogram.

Rasters from all 32 neurons responding to 100 Hz, 100 pA sinusoidal stimulation plus weak noise are shown in the top panel of **Figure 3D**. According to the firing rate histogram (constructed using 2.5 ms-wide bins), most spikes occurred between 2.5 and 5 ms after the start of each stimulus cycle (shaded bands). Though some cycles fail to evoke any spikes, most evoked phase locked spikes in 1-3 neurons, which represents <10% of the population being activated during any given stimulus cycle. A few hundred pyramidal neurons responding in this way – with sparse, phase locked spikes – can encode stimulus frequency despite noise (see below).

Next, we tested whether stimulus amplitude is encoded in the firing rate. For each neuron, firing rate for each stimulus amplitude was averaged across stimulus frequencies (**Fig. 4A**). Firing rate was significantly affected by both stimulus amplitude (*F*_1,28_ = 6.57, *p* = 0.016) and noise level (*F*_2,28_ = 44.83, *p* < 0.001) (2-way repeated measures ANOVA). Modulation of firing rate by stimulus amplitude was greatest in the presence of strong noise, but there was no significant interaction between the factors (*F*_2,28_ = 1.38, *p* = 0.260). Firing rate was also significantly affected by stimulus frequency (*F*_2,28_ = 21.49, *p* < 0.001) independent of noise level (*F*_4,28_ = 1.08, *p* = 0.370) (**Fig. 4B**). The increase in firing rate with increasing stimulus amplitude was consistent with expectations, but the decrease in firing rate with increasing stimulus frequency was unexpected and could compromise rate-based coding of stimulus intensity. Decreased responsiveness at higher stimulus frequencies is consistent with the low-pass filtering due to passive membrane properties (**Fig. 4C**) but the effects of the spike initiation process must also be taken into account (see Discussion).

**Figure 4.**
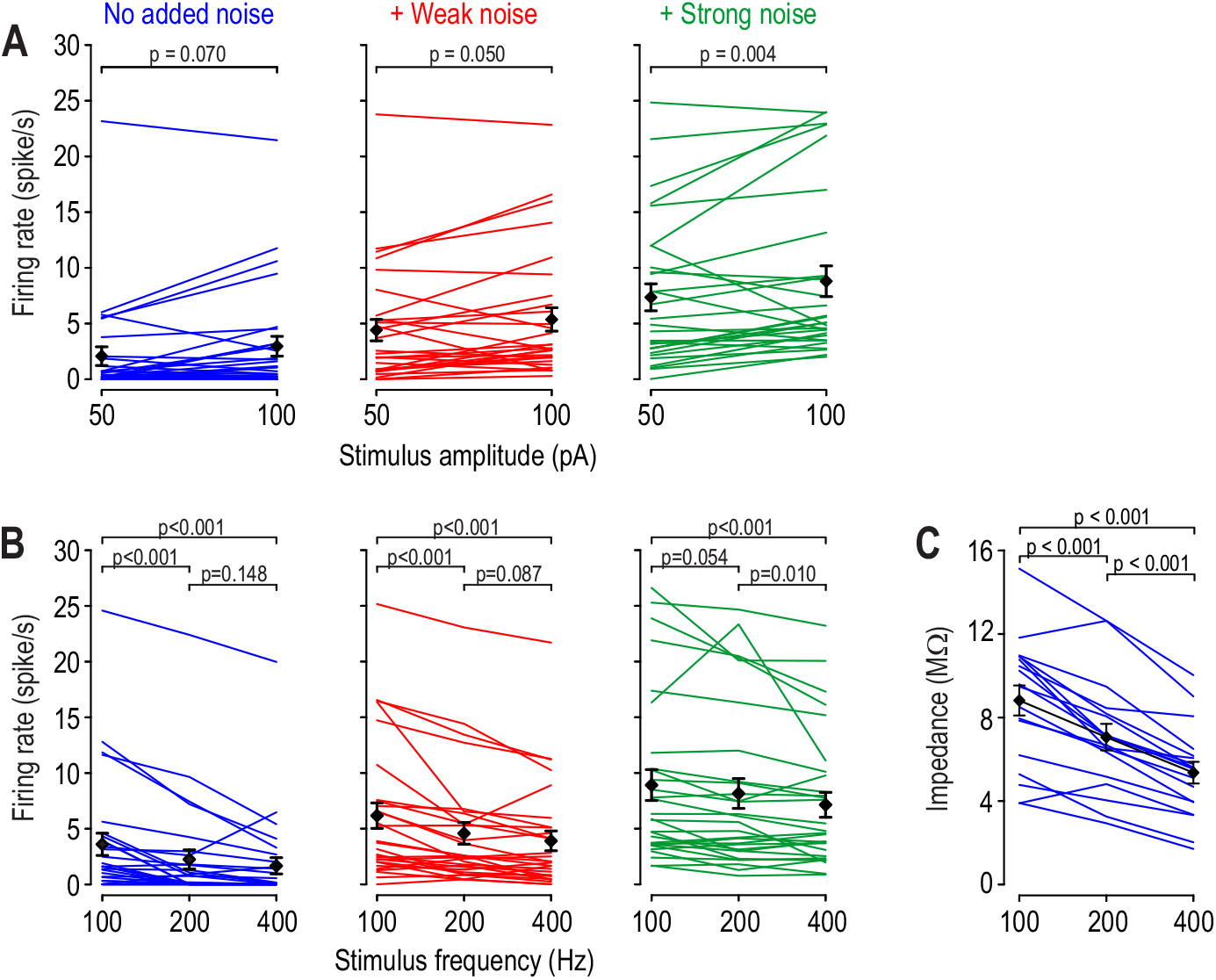
Firing rate is modulated by the amplitude and frequency of sinusoidal input. **(A)** Modulation of firing rate by stimulus amplitude. For each neuron, firing rate was averaged across all stimulus frequencies for each stimulus amplitude. Group averages (±SEM) are shown in black. Firing rate was affected by stimulus amplitude (*F*_1,28_ = 6.57, *p* = 0.016) and noise level (*F*_2,28_ = 44.83, *p* < 0.001) but there was no interaction (*F*_2,28_ = 1.38, *p* = 0.260; two-way repeated measures ANOVA). *P* values on graphs show results of post-hoc Student-Newman-Keuls tests. **(B)** Modulation of firing rate by stimulus frequency. Firing rate was averaged across stimulus amplitudes. Firing rate was affected by stimulus frequency (*F*_2,28_ = 21.49, *p* < 0.001). **(C)** Impedance, measured from subthreshold membrane potential fluctuations, was significantly lower at higher stimulus frequencies (*F*_2,17_ = 6.57, *p* < 0.001; one-way repeated measures ANOVA). Data are from separate recordings from 18 neurons.

### Multiplexed coding of trains of synaptic-like input by pyramidal neurons

Though primary sensory neurons experience mechanical stimuli whose amplitude (force) can vary sinusoidally in time, the input received by cortical neurons is re-shaped by the processing carried out by intervening neurons and synapses. Specifically, cortical neurons receive synaptic input triggered by volleys of spikes arriving near-synchronously at presynaptic terminals. With this in mind, we simulated trains of synaptic input by convolving a pulse train (100, 200 or 400 pulses/s) with a Gaussian kernel with a standard deviation of 0.5 ms (to account for spike jitter within each volley) and a synaptic waveform with a fast exponential rise and slower exponential decay (see inset on Fig. 5A). Virtual background synaptic input was unchanged. Like for sinusoidal input (see Fig. 3), synaptic trains evoked precisely timed spikes (**Fig. 5A**). Spike latencies were consistently short because the synaptic waveform kinetics are invariant, unlike for sine waves (where d*I*_stim_/d*t* increases with stimulus frequency). Spikes driven by synaptic trains remained precisely timed despite noise (**Fig. 5B**), as required for temporal coding of stimulus frequency.

**Figure 5.**
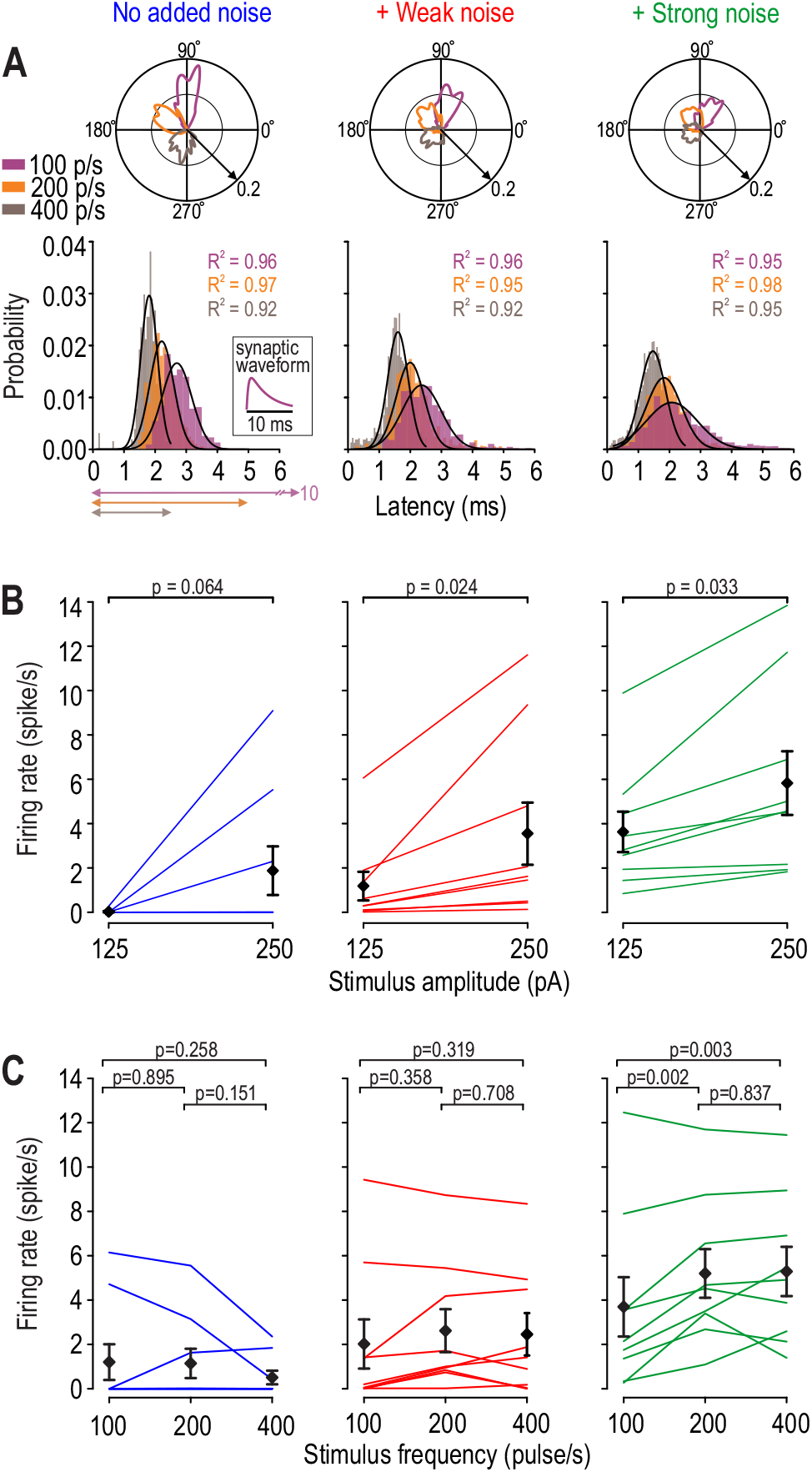
Trains of synaptic-like input drive phase locked spiking at a rate modulated by stimulus amplitude. **(A)** Distribution of spike times relative to stimulus phase (polar plots) and time from pulse onset (histograms). Data were aggregated across all neurons (*n* = 9) and stimulus amplitudes, and were fit with a Gaussian curve (black); coefficients of determination (*R*^2^) are reported on graphs. Inset shows the synaptic waveform. **(B)** Mean spike latency plotted against jitter for each combination of stimulus frequency, stimulus amplitude and noise level. Data are not shown for the 125 pA signal and no added noise condition because most neurons fired no spikes. Modulation of firing rate by stimulus amplitude. For each neuron, firing rate was averaged across stimulus frequencies. Group average firing rates (±SEM) are shown in black. Firing rate was affected by stimulus amplitude (*F*_1,8_ = 6.54, *p* = 0.034) and noise level (*F*_2,8_ = 32.75, *p* < 0.001) but there was no interaction (*F*_2,28_ = 0.475, *p* = 0.630) (two-way repeated measures ANOVA). *P* values on graphs show results of post-hoc Student-Newman-Keuls tests. Modulation of firing rate by stimulus frequency. Firing rate was averaged across stimulus amplitudes. Firing rate was not affected by stimulus frequency (*F*_2,8_ = 1.766, *p* = 0.203) but stimulus frequency interacted with noise level (*F*_4,8_ = 9.398, *p* < 0.001).

**Figure 5C** demonstrates that firing rate was significantly affected by stimulus amplitude (*F*_1,8_ = 6.54, *p* = 0.034) and noise level (*F*_2,8_ = 32.75, *p* < 0.001) (2-way repeated measures ANOVA), but there was no significant interaction between these factors (*F*_2,8_ = 0.475, *p* = 0.630). Nor was firing rate significantly affected by stimulus frequency (*F*_2,8_ = 1.766, *p* = 0.203) though there was a significant interaction between stimulus frequency and noise level (*F*_4,8_ = 9.40, *p* < 0.001) (**Fig. 5D**). This interaction likely arose because the synaptic waveform is longer than the interpulse interval (especially for high stimulus frequencies), which results in waveforms summating. The resulting sustained depolarization might be expected to increase spiking, but it also enhances adaptation via M-type K^+^ channel activation and Na^+^ channel inactivation, especially in the high-conductance state (Fernandez and White, 2010; Prescott et al., 2006). This sort of adaptation reduces spiking driven by fast voltage fluctuations (in the presence of noise) less than it reduces spiking driven by sustained depolarization (Fernandez et al., 2011; Prescott et al., 2008), which is consistent with results in Figure 5C. But if presynaptic neurons skip more cycles as stimulus frequency increases (see Figure 1), then fewer presynaptic neurons should fire on each stimulus cycle and the amplitude of synaptic input should decrease; this was not accounted for in the virtual synaptic input tested experimentally. Like Reyes et al. (1996), we incorporated a frequency-dependent decrease in synapse amplitude into a final set of simulations to explore this.

### Discrimination of stimulus amplitude and frequency in an AdEx model

Having established that pyramidal neurons, like the AdEx model, respond to periodic input with phase locked spiking that is resilient to physiological levels of noise, we returned to the AdEx model to quantify its ability to multiplex. Notably, many more stimulus parameter combinations can be tested in a model than can be tested with electrophysiological recordings of limited duration. Receiver operating characteristic (ROC) analysis was used to assess how well differences in stimulus amplitude or frequency could be discriminated based on differences in the rate or timing of spikes driven by sinusoidal input (**Fig. 6A**) or synaptic trains (**Fig. 6B**). For these simulations, synaptic input was modeled as an excitatory conductance (rather than current) and the amplitude of the synaptic waveform was scaled by stimulus frequency (see Methods), thus creating a more realistic input signal. For both types of input, differences in stimulus amplitude were discriminated from differences in spike rate (top panels) whereas differences in stimulus frequency were discriminated from differences in spike intervals (bottom panels). Frequency discrimination deteriorated for stimulus frequencies >400 Hz, especially for synaptic trains. Data in Figures 6A and B represent a single noise level, but testing with synaptic trains was repeated for several noise levels (**Fig. 6C**). Remarkably, discrimination of both stimulus amplitude and frequency was optimal near noise levels observed in the intact brain.

**Figure 6.**
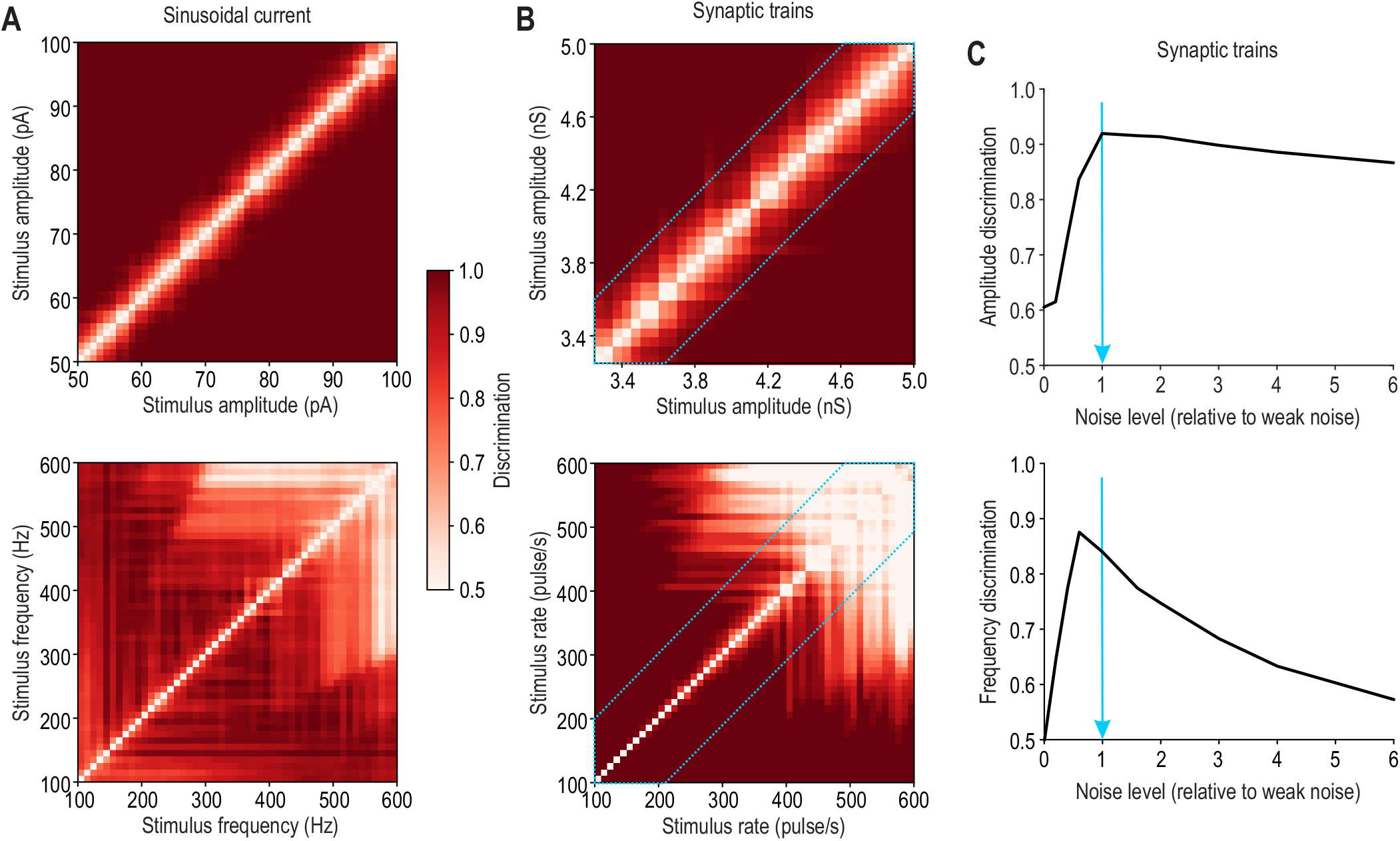
Discrimination of differences in stimulus amplitude and frequency in AdEx model based on spike rate and timing, respectively. Discrimination was assessed by ROC analysis and is reported here as area under the ROC curve (color). Discrimination of stimulus amplitude (top) and frequency (bottom) for sinusoidal current in high noise **(A)** and synaptic trains in low noise **(B)**. For each amplitude pair (top), 50 different randomly chosen frequencies were considered and the average discrimination is plotted, and vice versa for each frequency pair (bottom). In **B**, the area outlined in blue shows discrimination values that were averaged to produce a single discrimination value at each noise level in **C. (C)** Discrimination based on synaptic trains (as in **B**) was repeated for multiple noise levels expressed relative to “weak noise”, which corresponds our best estimate of noise in the intact cortex (blue arrow).

## DISCUSSION

Our results show that pyramidal neurons in S1 can use the rate and timing of their spikes to simultaneously represent (multiplex) the intensity and frequency of simulated vibrotactile inputs not despite noise, but because of it. Specifically, we found that L2/3 pyramidal neurons respond to high-frequency (up to 400 Hz) inputs with phase locked spikes under realistically noisy conditions. The resilience of phase locking to noise is surprising, but is consistent with *in vivo* recordings from S1 showing precisely timed spikes during tactile stimulation (see below). Our results argue that noise affects the probability (reliability) more than the timing (precision) of spikes evoked by small-amplitude, fast-onset inputs; in other words, reliability is more sensitive to physiological noise than precision under certain stimulus conditions. Moreover, when inputs repeat at an interval shorter than the neuron’s refractory period, the neuron responds intermittently, skipping many of its inputs. Noise causes inputs to be skipped irregularly, resulting in different neurons firing on different stimulus cycles so that the population as a whole can entrain to high-frequency stimuli. Firing rate encodes stimulus intensity because fewer cycles are skipped with increasing intensity. Independent modulation of spike probability and timing allows for independent rate- and time-based representations to be formed.

Vibrotactile stimuli evoke precisely timed spikes in primary afferents (Deschênes et al., 2003; Mackevicius et al., 2012; Pruszynski and Johansson, 2014; Weber et al., 2013), producing volleys of synchronized spikes that are relayed to the cortex (Bruno, 2011). Synchronized presynaptic spiking produces rapid postsynaptic depolarization (Rodriguez-Molina et al., 2007; Wang et al., 2010). The role of input synchrony is central to debates about coding strategy and operating mode, namely coincidence detection vs. integration (Abeles, 1982; London et al., 2010; Ratté et al., 2013; Shadlen and Newsome, 1998; Softky and Koch, 1993; Stevens and Zador, 1998). London et al. (2010) concluded that noisy conditions in cortex are incompatible with temporal coding *unless* there are fast depolarizing events. Such events clearly occur under certain stimulus conditions (DeWeese and Zador, 2006), including in S1 pyramidal neurons during mechanical stimulation (Crochet et al., 2011) due to stimulus kinetics or frictional interactions between the stimulus and the skin or whisker (*e*.*g*. slip-stick events) (Jadhav et al., 2009; Schwarz, 2016; Wolfe et al., 2008). Crochet et al. (2011) further observed that voltage deflections were strongly correlated across nearby neurons, consistent with neurons receiving common input, which is important for population coding since each neuron has a low (∼0.2) probability of spiking during each stimulus (Crochet et al., 2011). This is consistent with other data showing unreliable yet precisely timed spikes (Gabernet et al., 2005). That combination suggests that synaptic depolarization is abrupt (to produce well timed spikes) but relatively small (lest spikes occur more reliably), consistent with the precisely timed yet unreliable spikes we observed *in vitro*.

Past *in vitro* studies have shown that rapidly fluctuating inputs evoke spikes more reliably and with greater temporal precision than constant or slowly fluctuating inputs (Bryant and Segundo, 1976; Mainen and Sejnowski, 1995; Nowak et al., 1997; Rodriguez-Molina et al., 2007; Schreiber et al., 2009; Suter and Jaeger, 2004; Tiesinga et al., 2008). Rodriguez-Molina et al. (2007) found that only inputs with fast onset and decay kinetics evoked precisely timed spikes in the presence of background noise, and Khubieh et al. (2016) showed that noise can modulate spike probability without disrupting spike timing by randomizing the membrane potential prior to abrupt, signal-evoked depolarization. In the mean-driven regime (where spikes result from sustained suprathreshold depolarization), spiking is driven most reliably by input modulated at the same frequency as the ongoing firing rate, whereas in the fluctuation-driven regime the optimal input frequency is dictated by subthreshold resonance properties (Richardson et al., 2003; Schreiber et al., 2009). Non-optimal input frequencies reduce precision in the former condition and reliability in the latter (Schreiber et al., 2009). Our test conditions correspond to the fluctuation-driven regime.

We stimulated L2/3 pyramidal neurons by injecting current that varied sinusoidally or like a train of synaptic inputs. The latter more accurately reflects the input normally received by cortical neurons during vibrotactile stimulation, which evokes volleys of synchronized spikes in upstream neurons (see above). But direct stimulation by current injection excludes circuit-level processes like feedforward inhibition and synaptic processes like short term depression and facilitation that would normally contribute to shaping the synaptic input (Gabernet et al., 2005; Wright et al., 2021). Nonetheless, the periodic stimuli we used match the frequency and reasonably approximate the amplitude of synaptic inputs received by L2/3 neurons during vibrotactile stimulation (Crochet et al., 2011) and were sufficient to test whether pyramidal neurons generate precisely timed spikes given their intrinsic properties and noisy operating conditions. Early *in vitro* studies found that cortical neurons fire less as the frequency of sinusoidally modulated current is increased (Brumberg and Gutkin, 2007; Carandini et al., 1996), consistent with the low impedance associated with high frequencies. But subsequent work has emphasized that rapid spike initiation kinetics enable neurons to respond to frequencies much higher than predicted from passive membrane properties (Broicher et al., 2012; Eyal et al., 2014; Fourcaud-Trocmé et al., 2003; Higgs and Spain, 2009; Ilin et al., 2013; Kondgen et al., 2008; Tchumatchenko et al., 2011; Wei and Wolf, 2011). Our cells outperformed expectations based on simulations of rat L2/3 neurons (Eyal et al., 2014). Notably, the high-conductance state likely improves performance not only by reducing the membrane time constant, but by also speeding up spike initiation by electronically isolating the soma from the axon initial segment (Eyal et al., 2014).

By adding virtual synaptic input using dynamic clamp, we reproduced the noisy, high-conductance state normally experienced by L2/3 pyramidal neurons *in vivo* (Fernandez et al., 2018) and also tested stronger- and weaker-than-physiological levels of noise. Our results show that noise induces irregular skipping by making spike initiation more probabilistic. Inputs are subthreshold shortly after each spike because of the afterhyperpolarization (AHP), but they eventually reach threshold as the AHP wanes; noise can advance or delay that threshold-crossing by a few stimulus cycles, thus revealing the fundamental stimulus period. This reflects the combined effects of a slow drift in membrane potential due to the AHP and membrane potential randomization due to noise, as per Khubieh et. al (2016) (see above). The mechanism is less like stochastic resonance and more akin to dithering, or smoothing of the threshold nonlinearity, which can mitigate quantization artifacts (McDonnell et al., 2008). Larger inputs overcome the AHP faster, resulting in fewer skipped inputs, thus enabling stimulus intensity to modulate firing rate. In this scenario, stimulus intensity affects the amplitude but not the kinetics of each input. Noise plus small-but- controllable-amplitude inputs with consistently fast kinetics (due to presynaptic synchrony) allows for postsynaptic spiking with modulatable rate but precise timing.

Several *in vivo* studies have already established that at least some S1 neurons – depending on their afferent input (Saal et al., 2015) – respond to vibrotactile stimuli with precisely times spikes (Allitt et al., 2017; Arabzadeh et al., 2005; Ewert et al., 2015; Ewert et al., 2008; Harvey et al., 2013; Jadhav et al., 2009; Lieber and Bensmaia, 2019). The rate of those spikes can also carry information; for instance, Jadhav et al. (2009) found that firing rate and synchrony both increase with roughness, but synchrony varied more consistently than rate for subtle differences in texture. Lieber and Bensmaia (2019) found that variations in rate were sufficient for near-perfect discrimination of textures, suggesting the phase locked spiking they observed in some neurons is unimportant or relevant only to resolve subtler differences. Whether spike timing in S1 affects perception is contentious. Harvey et al. (2013) and Zuo et al. (2015) found that spike rate and timing both contributed to psychometric performance whereas Romo’s group – focusing on lower frequencies mind you – concluded that performance was accounted for by spike rate (Hernández et al., 2000; Luna et al., 2005; Salinas et al., 2000). That said, perceptual choice is more accurately reflected by activity in higher-order areas (Alvarez et al., 2015; de Lafuente and Romo, 2005; Kwon et al., 2016) where spike timing is poor (Harvey et al., 2013; Salinas et al., 2000). Even if time-based representations are eventually converted to rate-based ones, precisely timed spikes may play an important role in getting information about stimulus frequency to S1.

Data presented here establish that pyramidal neurons in S1 can support rate and temporal codes not despite noise, but because of it. The ability of cortical neurons to respond to fast depolarizing input with precisely timed yet unreliable spikes allows for phase locking and irregular skipping, respectively. That combination is critical for temporal coding and boils down to different factors determining *when* a spike will occur, and *if* it will occur. Modulation of skipping by stimulus intensity results in rate coding. Together these conditions allow the same spikes to encode the frequency and amplitude of periodic inputs using spike intervals (timing) and rate, respectively.

## MATERIALS AND METHODS

### Ethics Statement

All procedures were approved by the Animal Care Committee at The Hospital for Sick Children.

### Slice preparation

Adult male and female mice with a c57bl/6 background (Jackson Laboratory, Bar Harbor, ME) aged 6-8 weeks were deeply anesthetized with isoflurane and decapitated. The brain was rapidly removed from the skull and immersed in the ice-cold oxygenated (95% O_2_ and 5% CO_2_) sucrose-substituted artificial cerebrospinal fluid (aCSF) containing (in mM) 252 sucrose, 2.5 KCl, 2 CaCl_2_, 2 MgCl_2_, 10 D-glucose, 26 NaHCO_3_, 1.25 NaH_2_PO_4_, and 5 kynurenic acid. Coronal brain slices (300-400 µm) were prepared using a VT-1000S microtome (Leica, Wetzlar, Germany) and kept in oxygenated sucrose-substituted aCSF for 30 minutes at room temperature before being transferred to regular oxygenated aCSF (126 mM NaCl instead of sucrose and without kynurenic acid) and kept at room temperature until recording. Slices were transferred to a recording chamber constantly perfused with oxygenated aCSF heated to 31 ± 1 °C and viewed with an Examiner.A1 microscope (Zeiss, Oberkochen, Germany) using an IR-1000 Infrared CCD camera (Dage-MTI, Michigan City, IN).

### Patch clamp recordings

Pyramidal neurons in layers 2/3 of S1 were recorded in the whole-cell configuration with >70% series resistance compensation using an Axopatch 200B amplifier (Molecular Devices, Sunnyvale, CA). The pipette solution contained (in mM) 125 KMeSO_4_, 5 KCl, 10 HEPES, 2 MgCl_2_, 4 ATP, 0.4 GTP (Sigma-Aldrich); pH was adjusted to 7.2-7.3 with KOH. Reported values of membrane potential are corrected for a liquid junction potential of 9 mV. Data were low-pass filtered at 2 kHz and digitized at 20 kHz using a Power1401 computer interface and Signal 6.02 software (Cambridge Electronic Design, Cambridge, UK). Fast synaptic transmission was blocked via bath application of (in µM) 10 CNQX (Sigma-Aldrich), 40 D-APV (Tocris Bioscience), and 10 bicuculline (Sigma-Aldrich). Resting membrane potential, cell capacitance, and input resistance were intermittently re-checked during each recording and any neuron experiencing a large (>20%) change in any criterion was excluded from analysis.

### Periodic signals

Two types of periodic signals were tested experimentally. Both were applied by current injection. The first was a sine wave tested at three frequencies (100, 200, and 400 Hz) and two amplitudes (50 pA, 100 pA). The second was a synaptic pulse train in which pulses (at 100, 200, 400 pulse/s) were convolved with a Gaussian kernel with 0.5 ms standard deviation and with a synaptic waveform described by the Exp2Syn mechanism, namely

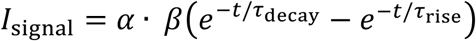

with τ_rise_ = 0.4 ms and τ_decay_ = 4 ms. The waveform’s peak is normalized to 1 by β so that the final amplitude corresponds to *α* = 125 or 250 pA. A DC component was adjusted for each neuron to evoke a similar initial firing rate.

### Virtual synaptic input (noise)

Background (random) excitatory and inhibitory synaptic input was applied using the dynamic clamp capabilities of Signal 6.02, where

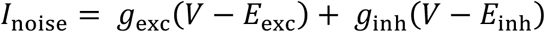

A junction potential error was added to reversal potentials (*E*_exc_ = 0 mV → +9 mV and *E*_inh_ = −70 mV → -61 mV) since the correction (see above) was only applied to voltages post-recording. Noisy fluctuations in excitatory and inhibitory conductance (*g*_exc_ and *g*_inh_) were modeled using separate Ornstein-Uhlenbeck processes where

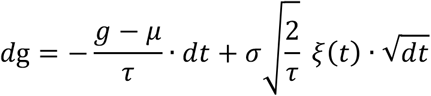

and *ξ* is a random number drawn from a Gaussian distribution with 0 average and unit variance, while τ_exc_ = 3 mS, τ_inh_ = 10 ms, *μ*_exc_ = 1 nS, *μ*_inh_ = 4 nS, σ_exc_ = 0, 0.25 or 0.5 nS and σ_inh_ = 0, 0.625 or 1.25 nS (for no noise, weak noise, and strong noise, respectively). This approach allowed us to separate the effects of noise from the effects of increased membrane conductance (Prescott and De Koninck, 2009), since background synaptic input increases both (Destexhe and Paré, 1999; Destexhe et al., 2003). With scaling factor 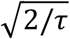, the standard deviation of the conductance corresponds to the value of σ prior to rectification. Conductances were rectified so that only positive values were applied. Weak noise (σ_exc_ = 0.25 nS and σ_inh_ = 0.625 nS) evoked membrane potential fluctuations of 2-2.5 mV, consistent with *in vivo* recordings from L2/3 pyramidal neurons in S1 (Fernandez et al., 2018).

### AdEx Model

Simulations were conducted using a modified AdEX model (Gerstner et al., 2014; Naud et al., 2008) according to the following equations

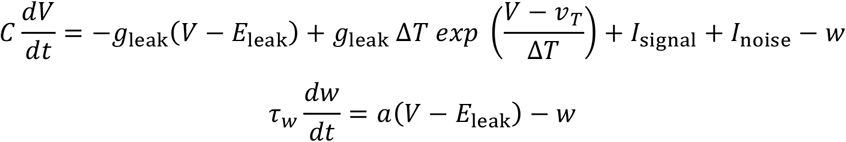

with threshold conditions such that at if *V* > *V*_th_, then *V* → *V*_reset_ *and w* → *w* + *b*. Parameters were: *C* = 80 pF, *g*_leak_ = 1 nS, *E*_leak_ = -65 mV, *a* = 1.3 nA, *b* = 0.25 nA, *v*_T_ = -45 mV, ΔT = 2 mV, *v*_reset_ = -70 mV, *τ*_w_ = 18 ms. A set of exploratory simulations were conducted before experiments to test initial predictions. After experiments, simulations were repeated with model parameters adjusted to more accurately reproduce typical experimental values including a capacitance of ∼80 pF, a resting membrane potential of ∼-65 mV, and a ∼70% drop input resistance upon introduction of the background synaptic input, similar to data from L2/3 pyramidal neurons recorded *in vivo* from S1 cortex (Fernandez et al., 2018). Results of the re-tuned model are reported.

Noise (*I*_noise_) was applied as rectified noisy excitatory and inhibitory conductances modeled as independent Ornstein-Uhlenbeck processes using the same parameters as in dynamic clamp experiments (see above). Signal (*I*_signal_) was either a sinusoidal current or synaptic input constructed as described above for experiments, but applied as a conductance. In the latter case,

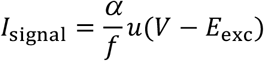

where *u* is the synaptic pulse train constructed as above, but where the scaling factor *α* is divided by the stimulus frequency *f* to produce a smaller synaptic input to account for the increased skipping that occurs at high stimulus frequencies. Specifically, if fewer presynaptic neurons fire on each stimulus cycle as stimulus frequency increases, then the amplitude of each synaptic input should decrease.

All simulations were performed in Brian2 (Stimberg et al., 2019) with the Euler-Maruyama algorithm with fixed time step of 10 µs. Model state (noise level, impedance, and stimulation type) was controlled with custom state control wrappers around Brian2. All code will be made freely available online upon publication.

### ROC Analysis

To assess how well stimulus amplitude and frequency were encoded, the AdEx model was tested with different combinations of stimulus amplitude and frequency. Receiver operating characteristic (ROC) analysis was then used to quantify discrimination of stimulus amplitudes based on spike rate or discrimination of stimulus frequencies based on spike timing. Discriminability is reported as area under the ROC curve. To decode spike timing, ISI distributions were computed. Since noise causes spikes to occur at integer multiples of the stimulus period, we expect to see peaks at

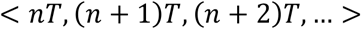

Therefore, the interval between peaks on the ISI histogram was measured and compared between stimulus frequencies using ROC analysis. For Figure 6C, area under the ROC curve is summarized at each noise level as the average within a strip along the diagonal for plots like those shown in Figure 6A and B. By focusing on this strip, we avoid overinflating the discriminability metric by including comparison of widely different stimulus amplitudes or frequencies (e.g. top left or bottom right corners of Fig. 6A, B).

## ACKNOWLEDGEMENTS

This work was supported by an Ontario Trillium Scholarship, Milligan Graduate Fellowship, Loo Geok Eng Foundation Scholarship and Vanier Scholarship to MAK; by a Restracomp fellowship to AS; by an NSERC University of Toronto Excellence Award to DS; and by a Canadian Institutes of Health Research Foundation Grant to SAP. We also thank Milad Lankarany for assistance with data analysis and stimulus construction, and Stéphanie Ratté for assistance with experiments and feedback on the manuscript.

## AUTHOR CONTRIBUTIONS

SAP conceived the study. DS and AS conducted simulations and analyzed data. MAK conducted experiments and analyzed data. MAK and SAP wrote the final paper with input from DS and AS.

